# Detection of ATTR aggregates in plasma of polyneuropathic ATTR-V30M amyloidosis patients

**DOI:** 10.1101/2024.05.13.593968

**Authors:** Rose Pedretti, Lanie Wang, Justin L. Grodin, Ahmad Masri, Jeffery Kelly, Lorena Saelices

## Abstract

ATTR amyloidosis is caused by the deposition of transthyretin amyloid fibrils in tissues often leading to organ failure and death. The clinical spectrum of this disease is highly diverse and dependent on many factors including the presence or absence of mutations within the transthyretin protein and/or an individual’s ancestry. The phenotypic variability of ATTR amyloidosis makes it difficult to diagnose, delaying treatment and worsening patient prognosis. Our lab has recently developed a peptide probe that detects transthyretin aggregates in plasma of ATTR amyloidosis patients with cardiomyopathy but has not been tested in plasma from polyneuropathic patients. Here we evaluate our probe in a cohort of Portuguese patients carrying the ATTR-V30M mutation and having no cardiac phenotype. We found that we could indeed detect aggregates in their plasma, and there appeared to be no relationship between the presence of aggregates and patient age or gender. Our work has broad implications on the pathobiology of ATTR amyloidosis and contribute to the validation of our probe as a novel detection tool for this disease.

---

Transthyretin amyloidosis is a disease caused by the systemic deposition of amyloidogenic transthyretin (ATTR) fibrils. The clinical spectrum of ATTR amyloidosis is highly diverse and may depend on factors such as the presence or absence of mutations within the transthyretin protein and an individual’s ancestry. For example, ATTR amyloidosis resulting from the wild-type transthyretin protein is thought to result primarily in infiltrative cardiomyopathy (ATTR-CA) later in life, whereas the V30M mutation in the Portuguese population is associated with peripheral neuropathy (ATTR-PN) early in life^1,2^. Diagnosing ATTR amyloidosis is challenging due in part to the phenotypic variability of the disease.

Recent studies indicate that transthyretin may aggregate in the blood of ATTR amyloidosis patients^3-5^. The Kelly lab has developed a peptide probe after the β-strand B of transthyretin that detects non-native transthyretin (NNTTR) species in plasma of ATTR-PN patients with the V30M mutation^3^. These species are absent in those patients with cardiomyopathic or mixed phenotypes^3^. Our lab has also developed a <u>T</u>ransthyretin <u>A</u>ggregation <u>D</u>etection (TAD1) probe targeting β-strands F and H that detects aggregated transthyretin species in plasma of ATTR-CA patients and patients with mixed phenotypes of both cardiomyopathy and polyneuropathy^4,5^. TAD1, however, has not been validated using plasma samples from ATTR-PN patients yet. Thus, here we assess whether the TAD1 probe can detect similar aggregated species in patients with strictly polyneuropathic phenotypes.

Raw data, patient data, and analytical methods can be made available upon request. Human tissues and peptide probes cannot be made available because of legal constraints. The Office of the Human Research Protection Program at UTSW granted exemption from Internal Board review because specimens are anonymized. Experiments were conducted by the researchers blinded and patient information was revealed for analysis at the conclusion of the study.

We sought to test the TAD1 probe’s ability to bind aggregates in plasma of ATTR amyloidosis patients of varying clinical phenotype. We tested this using three distinct cohorts: ATTR-PN patients, ATTR-CA patients, and technical control plasma samples that do not bind TAD1 (Figure 1A). We subjected these cohorts to our TAD1 assay pipeline, where TAD1 is incubated with plasma fixed on nitrocellulose membrane, followed by washing steps and quantifying aggregates in the samples through TAD1 fluorescence intensity (Figure 1B). We found that TAD1 detects aggregates in ATTR amyloidosis plasma regardless of whether the patient suffers from a predominantly cardiac or neuropathic phenotype (Figure 1C). We found no significant correlation between TAD1 signal and the patients’ gender (Figure 1D) or age (Figure 1E). Even though the ATTR-PN cohort is much younger than the ATTR-CA cohort, their TAD1 signals are comparable.

**Figure 1.**
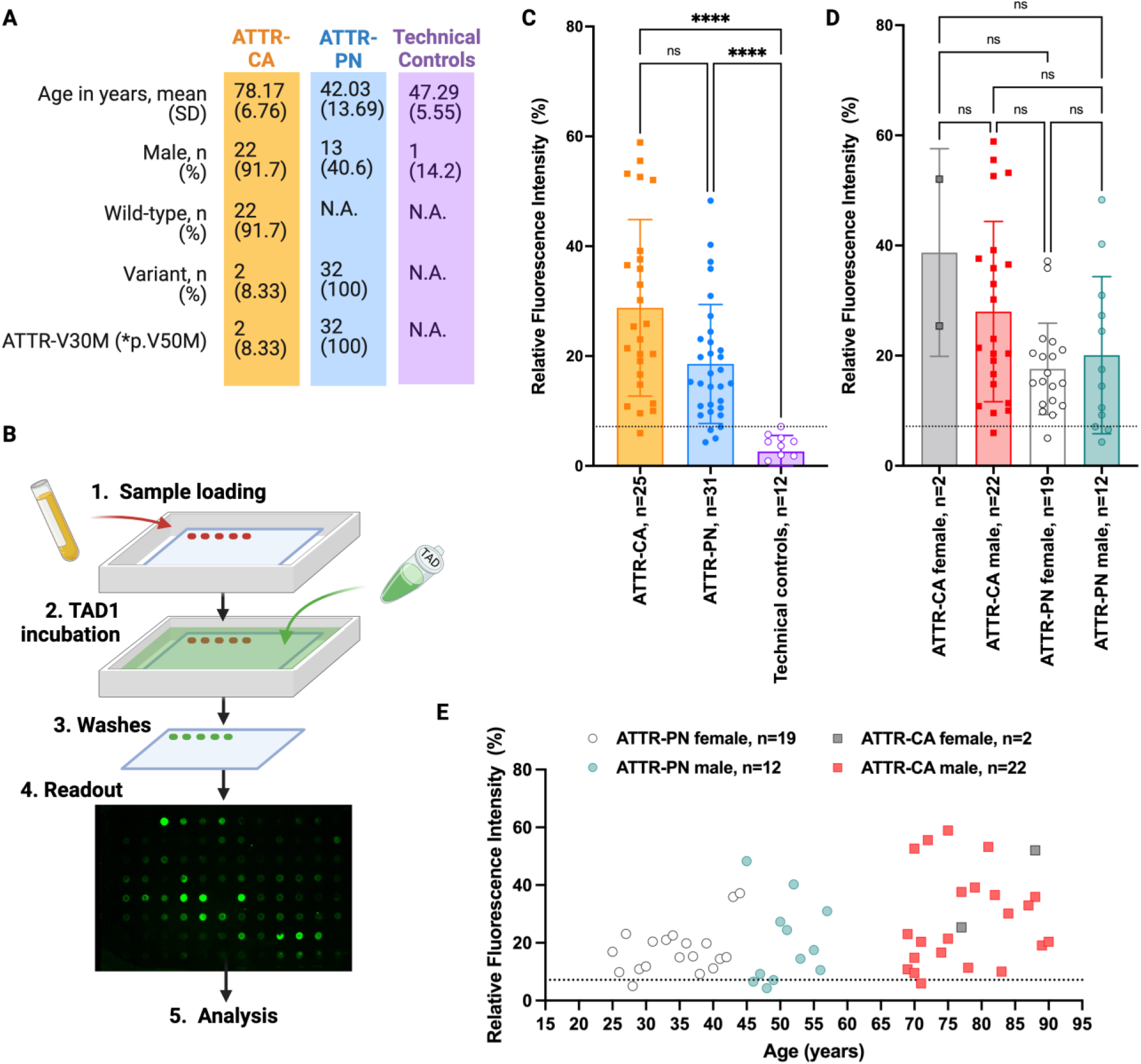
TAD1 detects ATTR aggregates in plasma of neuropathic ATTR-V30M patients. **A**. Characteristics of plasma samples used for the study. **B**. Schematic of the protocol used for analyzing aggregates in plasma. 30 µL of plasma samples are loaded onto nitrocellulose membrane then incubated with 5 µM of our <u>T</u>ransthyretin <u>A</u>ggregation <u>D</u>etector 1 (TAD1) probe overnight. Excess unbound TAD1 is washed off the membrane through three ten-minute washes with 10% tris buffered saline, then the level of ATTR aggregates in samples are measured through excitation of TAD1 at 472 nm and reading emission at 513 nm. Fluorescence intensity is quantified using ImageJ **(C-E)**. Signal is normalized to membrane as 0% and 0.5 µg *ex vivo* ATTRwt fibrils as 100%. **C**. TAD1 detects ATTR aggregates in both cardiomyopathic (ATTR-CA) and polyneuropathic (ATTR-PN) ATTR amyloidosis patients. Outliers were removed using a Grubbs test of the normalized dataset. A non-parametric Mann-Whitney *t*-test was used to establish significant differences between each pair of groups (ns, not significant; **** p <0.0001). Dashed line denotes the highest value of technical controls that do not bind TAD1. **D**. There is no correlation between the gender of the patient and TAD1 signal. Statistical analysis was performed as in **C. E**. There appears to be no relationship between age and TAD1 signal for both cohorts assessed.

Our findings may have important implications on the biology of ATTR amyloidosis as well as the development of clinical tools to detect and treat disease. Combined with previous studies, our results suggest that in ATTR-CA patients, there is only one type of prefibrillar aggregated species in plasma, recognized by TAD1. However, it appears that ATTR-PN patients may have multiple types of aggregated ATTR species, those that are detected by the NNTTR detection assay^3^ and those detected by TAD1^4,5^. This may or may not relate to the phenotypic variability observed in ATTR amyloidosis patients and warrants further study. In combination, the probes developed by our lab and the Kelly lab could have potential to be used as a blood-based diagnostic tool to differentiate between ATTR-CA and ATTR-PN.

There are limitations to this study that are of importance to note. These limitations include a small sample size that does not have the power to adjust for confounding variables. These samples were obtained through convenience sampling and needs independent validation with a larger sample size and a targeted experimental design.

In summary, the present study evaluates the presence of aggregated transthyretin species in plasma of neuropathic ATTR amyloidosis patients, similar to those found in cardiac ATTR amyloidosis patients^4,5^. We observed that patients with a predominantly neuropathic phenotype contain similar ATTR species in their plasma, which are detected by our novel TAD1 probe. These findings may have important implications on the biological and clinical aspects of ATTR amyloidosis.

## Notes

### Competing Interest Statement

RP and LS are inventors on a patent application (Provisional Patent Application 63/352,521) submitted by the University of Texas Southwestern Medical Center that covers the composition and structure-based diagnostic methods related to cardiac ATTR amyloidosis. JLG receives honoraria for scientific consulting for Alnylam, Eidos/BridgeBio, Intellia, Pfizer, Alexion, Astra-Zeneca, and Tenax Therapeutics and receives research funding from Pfizer, Eidos/BridgeBio, the Texas Health Resources Clinical Scholars fund, and the NHLBI. AM receives research funding from Pfizer, Ionis/Akcea, Attralus, and Cytokinetics. AM also receives fees from Cytokinetics, BMS, Eidos, Pfizer, Ionis, Lexicon, Alnylam, Attralus, Haya, Intellia, BioMarin, and Tenaya. LS receives honoraria for scientific consulting for Intellia and Attralus, and serves as a member of the advisory board of Alexion.

